# A sub-set of guanine- and cytosine-rich genes are actively transcribed at the nuclear Lamin B1 region

**DOI:** 10.1101/2023.10.28.564411

**Authors:** Gayan I. Balasooriya, Payal Naik, Tse-Luen Wee, David L. Spector

## Abstract

Chromatin organization in the mammalian cell nucleus plays a vital role in the regulation of gene expression. The lamina-associated domain at the inner nuclear membrane has been shown to harbor heterochromatin, while the nuclear interior has been shown to contain most of the euchromatin. Here, we show that a sub-set of actively transcribing genes, marked by RNA Pol II pSer2, are associated with Lamin B1 at the inner nuclear envelope in mouse embryonic stem cells (mESCs) and the number of genes proportionally increases upon *in vitro* differentiation of mESC to olfactory precursor cells. These nuclear periphery-associated actively transcribing genes primarily represent housekeeping genes, and their gene bodies are significantly enriched with guanine and cytosine compared to genes actively transcribed at the nuclear interior. We found the promoters of these gene’s to also be significantly enriched with guanine and to be predominantly regulated by zinc finger protein transcription factors. We provide evidence supporting the emerging notion that the Lamin B1 region is not solely transcriptionally silent.

## Introduction

Euchromatin and heterochromatin of mammalian nuclei are compartmentalized into independently distinguishable nuclear regions [1-4]. Early microscopic observations have shown that the heterochromatic histone signatures mark denser chromatin structures at the nuclear periphery (NP) and at internal dense chromatin domains, and the bulk of diffuse chromatin at the nuclear interior (NI) is marked by euchromatic histone signatures that are present on transcriptionally active genes [3, 4]. Furthermore, chromatin capture techniques (e.g., Hi-C and 5C) also provide evidence that chromatin in the nucleus is compartmentalized into two Hi-C-defined nuclear compartments, the transcriptionally active gene containing nuclear compartment A and the transcriptionally silent gene containing nuclear compartment B [5]. The nuclear lamina is primarily comprised of Lamin A/C and Lamin B, mainly Lamin B1 (LB1) [6, 7], as well as inner NE proteins such as Lap2, Emerin and Man1 [8]. The chromatin at the NP has been shown to interact with the nuclear lamina [9] and therefore, the nuclear lamina plays a vital role in chromatin organization and the regulation of gene expression in the cell nucleus.

Further, the Dam-ID technique, used to label chromatin interacting with LB1 (lamina-associated domains or LADs), has also shown that the B compartment is predominantly associated with LADs at the NP [10]. However, the nuclear pore regions lacking the lamina meshwork have been shown to “gate” transcriptionally active genes [11, 12] and thus, the NE does not represent a completely heterochromatic region [13-15]. Further, by performing structure-resolving image analysis for LB1, it has been shown that inactive (H3K27me3) and poised (H3K27ac) enhancers, both marked by H3K79me2, therefore having the potential to gain activation, are present at the LB1 low-abundant Hi-C defined A compartment at the NP [16].

It has been shown that when cells differentiate, the transcriptional identity and the composition of genes at the LB1 regions change [17, 18]. Further, the arrangement of euchromatin and heterochromatin in a nucleus can also change during cell specification; some genes move away from the LB1 region to be activated, and some move into the LB1 region to be activated [12, 19]. For example, most of the euchromatin in olfactory cells, the rod cells, and Lamin B receptor-null thymocytes is present at the NP [20-22]. However, the association of those transcriptionally active genes within the different sub-compartments (e.g., LB1, nuclear pores, LB1 adjacent) of the inner NE has generally not been investigated in detail.

To date, the presence of heterochromatic genes at the nuclear periphery (NP) was identified based on data generated from histone signatures, Hi-C, and Dam_ID methods [23-26]. To our knowledge, approaches specifically using transcriptional profiling of the LB1-associated genes based on RNA polymerase II (RNA Pol II) activity have not yet been investigated in detail genome-wide. Although transcriptional profiling by histone signatures provides the overall status of a gene’s transcription (e.g., active or silent), the different state’s of a gene’s transcription (pre-initiation, promoter-proximal pausing, elongation, termination) is not informative from such data.

However, occupancy by RNA Pol II and its phosphorylated derivatives at the different regions of a gene can provide more insight into its transcriptional status. Transcription can be explained as a step-by-step process: RNA Pol II recruitment, initiation, elongation, and termination. Although the phosphorylation of Ser5 at the C-terminal domain of the RNA Pol II large subunit defines the transcription initiation of a gene, a gene in such a state can represent transcription elongation (in ‘canonical’ transcription) or promoter-proximal pausing [27, 28]. However, phosphorylation at Ser2 (pSer2) at the C-terminus of RNA Pol II is consistent with transcriptional elongation and thus can be used as a molecular signature of actively transcribing genes. To our knowledge, the presence of RNA Pol II pSer2 association, and therefore actively transcribing genes, at the LB1 region, has not been investigated in depth. Although the bulk data from RNA Pol II pSer2-associated chromatin profiling using chromatin immunoprecipitation (ChIP) is available [29-31], those data are non-informative for distinguishing actively transcribed genes residing in different nuclear compartments.

Since the LB1 region at the NP has been shown to harbor a large amount of DNA [2] we focused on investigating the possibility of actively transcribing genes being localized with LB1 at the NP using immunofluorescence (IF) microscopy and sequential ChIP-seq (using LB1 and RNA Pol II pSer2 specific antibodies) assays. In mouse embryonic stem cells (mESCs) and *in vitro* mESC-derived neural progenitor cells (NPCs - single clone derived), we found that a subset of actively transcribing genes was associated with LB1 at the NP. Further, we demonstrated that upon mESC differentiation toward NPCs, the number of actively transcribing genes at the LB1 region was increased in our clonally derived “NPCs”. Using transcriptomic signatures and immunofluorescence labeling for Ascl1 (Globose basal cell signature), we showed that the *in vitro*-derived “NPCs” we used in our study were in fact olfactory progenitor cells (OPCs). Mature olfactory cells have a reverse nuclear organization whereby euchromatin is partially present at the NP and heterochromatin at the NI [22]. Therefore, the *in vitro*-derived “NPCs” are primed OPCs that are setting up the chromatin organization of mature olfactory cells. Moreover, we found that actively transcribing genes at the LB1 region are significantly enriched with guanine and cytosine and contain relatively lengthier gene bodies than those that are transcribed at the NI. Although we did not find these genes relevant to the cell identity of OPCs, they primarily represent housekeeping genes. Further, the promoters associated with these genes are also enriched with guanine and are known to be predominately regulated by zinc-finger transcription factor family proteins. In summary, we present evidence that actively transcribing genes exist at the LB1 compartment and they exhibit distinct molecular characteristics from genes transcribed at the NI.

## Materials and Methods

### Mouse ESC (mESC) culture and *in vitro* neural progenitor cell (NPC) derivation

Polymorphic F1 mESCs were cultured *in vitro* as previously described [32]. To derive NPCs from mESCs, male F1 mESCs were seeded on 0.1% gelatin-coated at 1×10^5^ cells per well in 6-well plates with 2.5 ml 2i medium. After one day of seeding, cells were washed three times with 1x PBS and 2 ml of cell differentiation medium (10% FBS (Millipore, Cat. ES-009-B, 1X L-Glutamine (Thermo Fisher, Cat. 25030081, 1X Non-Essential AA, (Millipore, Cat. TMS-001-C), 0.15 mM 2-Mercaptoethanol (Millipore, Cat. ES-007-E), 100 U/ml Penicillin-Streptomycin (Gibco, Cat. 15140-112) in Knockout DMEM (Gibco, Cat. 10829-018) added to each well and incubated at 37 ^ο^C, 5% CO2 for 24 hrs. Cells were then washed three times with 1xPBS and cultured with 2.5 ml cell differentiation medium supplemented with 2.5 µg/µl murine EGF (PeproTech Inc, Cat. 315-09) and 2.5 ug/ul murine Fgf2 (PeproTech Inc, Cat. 450-33). After 48 hrs, cells were dissociated by TrypLE Express Enzyme (Thermo Fisher, Cat. 12605028) and seeded at 1×10^5^ cells per well in 0.1% gelatin-coated 6-well plates in cell differentiation medium supplemented with EGF and Fgf2 for subsequent long-term NPC cultures. From the NPC cultures, we established a three single clone-derived NPC lines to be further used in our study.

### Immunofluorescence assay

Cells were cultured on ibidi u-Slide 8-well glass bottom chambers (ibidi GmbH, Cat. 80827). Cells were washed once with 1x PBS and fixed in 4% PFA for 15 min while gently rocking, and the PFA was removed and the cells rinsed with 1xPBS three times, 10 min for each wash. Cell permeabilization was performed using 0.3% PBSTx (Triton X-100 in 1xPBS) for 15 min at room temperature (RT) while gently rocking on a rocking platform. Cell permeabilization buffer was removed, and the cells were washed with 0.1% PBSTx three times (10 min for each wash). 200 µl of blocking buffer (3% BSA, 10% NDS, 0.1% PBSTx) was added per well and incubated for 1 hr at RT. 200 µl of blocking buffer containing primary antibodies was added to each well and incubated on a shaker. Primary antibodies were rinsed off, and the cells were then washed with 0.1% PBSTx four times (10 min for each wash). The secondary antibody diluted in 200 µl of the secondary antibody buffer (5% NDS in 0.1% PBSTx) was added to each well and incubated at RT on a slow rocker. After the incubation period, secondary antibodies were rinsed off, and then the cells were washed with 0.1% PBSTx three times and finally washed once with 1X PBS. DAPI diluted at 1:10,000 in 1xPBS was (200 µl) added to each well, incubated for another 5 min, and then washed three times with 1X PBS. 100 µl of ibidi mounting medium (ibidi GmbH, Cat. 50001) was added to each well, and the chamber slides were kept wrapped in aluminum foil at 4 °C until imaged. Primary antibodies used: Lamin_B1 (1:1,000) (Abcam, Cat. ab16048), RNA Pol II pSer2 (1:500) (ActiveMotif, Cat. 61084), Pax6 (1:500 – BioLegend, Cat. 901301), Nestin (1: 500 – Abcam, ab6142), △Np63 (1:500 - BioLegend, Cat. 699501), Ascl1 (1:500 - NOVUS Biologicals, Cat. NBP3-17687). Secondary antibodies used: Goat anti-rat (Thermo Fisher, Cat. A-11006), Donkey anti-mouse (Thermo Fisher, Cat. A-31570), Donkey anti-rabbit (Thermo Fisher, Cat. A-31573). Secondary antibodies were used at 1:1,000 dilution. Lamin_B1 and RNA Pol II pSer2 primary antibodies were incubated with cells for 45 min, and then the secondary antibodies were also incubated with cells for 45 min at RT on a slow rocker. All the other primary antibodies were incubated overnight at 4 °C, and then the cells were incubated with secondary antibodies for 1.5 hrs at RT on a slow rocker.

### Imaging

Cells were imaged using the Spinning Disk Laser Scanning Confocal Microscope (UltravVIEW Vox, PerkinElmer) with CSU-X1 scan head with 50 µm pinholes (Yokogawa) integrated in an inverted imaging platform and a Nikon (Nikon Japan, Plan Apo VC, 100x/1.40 oil, ∞/0.17 DIC 2N) lens. The platform incorporates diode-pumped solid-state (DPSS) lasers (Vortran VersaLase™) with an ORCA-Fusion BT scientific CMOS camera (C15440-20UP, Hamamatsu, Japan). The camera was set to 16-bit high-quality mode with no binning. The platform is integrated with Volocity image acquisition software (version 7.0.0, Quorum Technologies). For the RNA Pol ll pSer2 and LB1 IF experiment, sequential imaging was conducted under the DAPI channel (405nm laser) at 50% laser power, 300ms exposure time, GFP channel (488nm laser) at 55% laser power, 400ms exposure time, Cy5 channel (635nm laser) at 51% laser power, 400ms exposure time. Two X-Y plane images 3 μm apart were taken for each cell. Nascent RNA-FISH samples were imaged using the same settings as above for the GFP and Cy5 channels. The Z-slice of 0.3 μm Z-spacing was used by moving the ASI automated XYPZ stage (MS-2000, Applied Scientific Instrument) upwards, capturing multiple channels in each Z plane and collectively capturing images from each entire cell. For the histone marks and LB1 IF experiment, sequential imaging was conducted under the DAPI channel (405nm laser) at 17.3% laser power, 300ms exposure time, the AF488 channel (488nm laser) at 51.1% laser power, 800ms exposure time, and the AF568 channel (561nm laser) at 51.1% laser power, 800ms exposure time. Each 3D image stack was acquired with 0.3 µm spacing between Z-slices, collected in channel-first order, followed by the Z-plane using the ASI automated XYPZ stage (MS-2000, Applied Scientific Instruments).

### Image analysis

Image analysis was performed using ImageJ2 (Version: 2.24.0/1.54f) software. Prior to colocalization analysis, channels were split, and the background was subtracted using the Rolling ball radius method (setting at 50.0 pixels) and setting parameters for the Sliding paraboloid. The background corrected images were then used for colocalization analysis using the Coloc 2 plugin. Colocalization coefficiencies were calculated using Manders’ Correlation and Spearman’s Rank Correlation algorithms without setting ROI or mask for the objects. The average values of the thresholds from each analysis and standard error of means (SEM) were calculated and presented in the bar graphs.

### Immunoblotting

Immunoblotting was performed as in our previous manuscript [33] with different antibodies. Briefly, Tris-glycine SDS-polyacrylamide gels (5% Stacking gels - 30% acrylamide mix, 1.0 M Tris (pH6.8), 10% SDS, 10% ammonium persulfate, 0.1% TEMED in dH_2_O and 12% resolving gels - 30% acrylamide mix, 1.5 M Tris (pH8.8), 10% SDS, 10% ammonium persulfate, 0.1% TEMED in dH_2_O) were cast 1.00 mm thick and 8×10 cm gels were used for protein electrophoresis. Proteins were transferred from the gel in the ice-cold 1x transfer buffer onto Immobilon®-P Transfer membranes (Merck Millipore, Cat. IPVH00005). Transfer membranes were air-dried to dehydrate and stored until use. Membranes were rehydrated in 0.1% PBST (0.1% Tween20 in 1xPBS), changing the buffer several times until the membrane became wet before probing with antibodies. Rehydrated membranes were blocked in 5% blocking buffer containing skim milk (Sigma, Cat. 1153630500) in 0.1% PBST for 1 hr at room temperature and then incubated with primary antibodies diluted in blocking buffer with 5% milk overnight at 4 °C. Membranes were then washed in 0.1% PBST four times, changing the buffer every 10 minutes, and then incubated with secondary antibodies conjugated with horseradish peroxidase diluted in secondary antibody buffer (3% milk in 0.1% PBSTX) for 2 hrs at RT on a shaker. After the incubation, membranes were washed four times (10 minutes each) with 0.1% PBST buffer and, lastly, once with 1x PBS. The ECL (Thermo Fisher, Cat. No. PI32209) was added onto the membrane to cover the whole membrane and it was exposed to X-ray film (Amersham Hyperfilm™ ECL, GE Healthcare Limited, Cat. 28906836). Primary antibody used: Lamin_B1(1:1,000) (Abcam, Cat. ab16048), Secondary antibody: Donkey anti-Rabbit IgG, (H+L) Secondary Antibody, HRP (Thermo Fisher, Cat. 31458).

### Sequential Chromatin Immunoprecipitation

Cells were expanded in 10 cm dishes up to ∼80% confluency, and 1 x 10^7^ cells were used per IP. To be compatible with a sequential ChIP assay, we used a native (non-cross-linking) approach. Single cells obtained after dissociation were resuspended in 3,000 µl RIPA buffer (0.1% SDS, 10 mM EDTA, 50 mM Tris-HCl (pH 7.5), 1 mM PMFS, Protease inhibitor (Roche, Cat. 39802300)) and sonicated in 15 ml tubes containing ∼25 µl of sonication beads (diameter > 1 mm) using Bioruptor® pico (Diagenode, Cat. B01060010) for 15 cycles with 30 sec ON and 30 sec OFF for each cycle in ice-cold water. Sonicated samples were kept in ice until the sonication beads were settled to the bottom of the 15 ml tube, and then the supernatants were transferred into separate vials. Pre-washed (Once with 500 µl of 0.02X Tween20 in 1xPBS) 100 µl Dynabeads Protein G (Invitrogen, Cat. 10004D) was used per sample. Before incubating with antibodies, Protein G beads were blocked with 0.5% (w/v) BSA for 1 hr at RT and washed with 1 ml of 1X PBS three times. Beads were then resuspended in 500 µl of 0.02X Tween20 in 1xPBS and incubated with 8 µg of Lamin B1 (Abcam, Cat. ab16048) and 8 µg of IgG (Millipore, Cat.12-370) per sample for one hour on a rotator at 4 °C. Supernatant from each tube was removed by pipetting out while the tubes were on a magnetic rack. Then, the beads were washed once with 1 ml washing buffer (0.02X Tween20 in 1xPBS), and 1,000 µl from the sonicated samples were added to each tube containing Protein G beads incubated with Lamin_B1 antibody. Samples were incubated on a rotator at 4 °C overnight. After incubation, the supernatant was pipetted out while the tubes were on the magnetic rack, and then the beads were washed three times (5 min for each wash) on the rotator at 4 °C with 1 ml cold Re-ChIP buffer (2 mM EDTA, 500 mM NaCl, 0.1% SDS, 1% NP40). Beads were then rinsed once with 1x TE buffer, and the chromatin-Lamin B1 complexes were eluted from beads using 150 µl (per sample) Re-ChIP elution buffer (1x TE, 2% SDS, 10 mM DTT, protease inhibitor) by incubating the samples at 37 °C for 30 min on a rotor set at 800 rpm. Finally, chromatin-Lamin B1 complexes were eluted and diluted by adding 850 µl of ChIP dilution buffer (1.1 % TritonX-100, 1.2 mM EDTA, 16.7 mM Tris-HCl (pH 8.0), 167 mM NaCl, proteinase inhibitor) to use in Pol II pSer2 pull down.

Pre-blocked and washed (same as in the first step), 150 µl of Dynabeads Protein G beads were used for each sample in the second immunoprecipitation step. RNA Pol II pSer2 [34] and IgM (Invitrogen, Cat. 11E10-14-4752-82) isotype controls (2 µg from each) diluted in 150 µl of Re-ChIP buffer were incubated with Dynabeads Protein G for 2 hr at 4 °C on a rotator. The supernatants were pipetted out while the tubes remained on a magnetic rack and mixed with Lamin_B1 ChIP’ed samples (850 µl for each reaction sample). Samples were incubated at 4 °C overnight and then processed for chromatin isolation. Each sample was washed first (in 1 ml) with cold low-salt wash buffer (0.1% SDS, 1% Tritronx-100, 2 mM EDTA, 20 mM Tris-HCl (pH 8.0), 150 mM NaCl) and in cold High-Salt Buffer (0.1% SDS, 1% Tritron X-100, 2 mM EDTA, 20 mM Tris-HCl (pH 8.0), 500 mM NaCl) and in cold Li-Cl Buffer (0.25 M LiCl, 1% NP40, 1% DOC, 1 mM EDTA, 10 mM Tris-HCl (pH 8.0) and lastly with cold TE buffer (10 mM Tris-HCl (pH8.0), 2 mM EDTA). Washes were carried out 3X for 10 min each on a rotator set at 800 rpm. DNAs were isolated using a Genomic DNA Clean & Concentrator™ -25 Kit (Zymo Research, Cat. D4064).

### DNA sequencing

Paired-end libraries were prepared using Illumina TruSeq ChIP library preparation kit (Illumina, Cat. IP-202-1012) and sequenced (PE-150, mid output) using NextSeq500 sequencer at the Next Generation Sequencing Core Facility at Cold Spring Harbor Laboratory.

### ChIP-seq analysis

The MEA pipeline[35] installed in the CSHL supercomputing cluster was used for ChIP-seq analysis, as explained in our previous study [32]. In silico diploid C57BL/6J / CAST/EiJ genome was reconstructed using the SHAPEIT2 pipeline by combining the mm10 reference genome (mgp.v5.merged.snps_all.dbSNP142.vcf.gz) and genetic variants (mgp.v5.merged.indels.dbSNP142.normed.vcf.gz). We then aligned sequencing read files (.fastq) to the constructed in silico genome using Bowtie2 aligner (version 2.3.4.3). Generated ‘.bam’ files were then used in the ChIP-seq pipeline embedded in the SeqMonk Mapped Sequence Analysis tool from the Babraham Institute, UK. Peaks were called using the contig caller embedded in the SeqMonk software package with flowing parameters: readcount=/>10, fragment length=500 bp, using input as the control. ChIP peaks were visualized using Integrative Genomic Viewer (IGV)[36].

### Nick translated probe preparation and nascent RNA-FISH

Nascent RNA-FISH assay was performed as previously described [33]. Briefly, genomic DNA from mESCs was used as a template to PCR amplify the DNA probes used in nick translated probe preparation. Primers used for PCR amplification and used as a template: Zfhx3: F-TTTCGCACTCAAATGACCAA, R-CTCAAAAAGGTGGGCAGAAG, Fgfr2: F-TCCCTAACCCGTTTCTGTTG, R-CAGCATACATGGTGGGTCAG, Cntnap2: F-AACTAGGCAGGTCTGGAGCA, R-GCCAGTGGTACTCCAAAAGC, Lsamp: F-GGACCCAGGCATAAATGCTA, R-CACCAAAGGCAAGACCTTCT, Klf3: F-ACAAGCTTTCAGTGCACAGG, R-GAAGGTCTCTGGCAAGGACT, Rab33b-GGGCCTCTCATAGCACTAGG, R-CAATCCCCACTGTAAAGCGG, Et14: F-TCCTCCATTACCCCAGTGTC, R-GGTATGTTTCTGGGCTTCCA, Macf1: F-ACCGACACCAGCTTTTCACT, R-CCCAATGTATGGAGCAGGAT.

One microgram of DNA probe from PCR was used in each reaction. The reaction mixture comprised DNA fragments, 3.8 µL of labeled dUTP (0.2 mM), 2.5 µL of dTTP (0.1 mM), 10 µL of dNTP mix (0.1 mM), 5.0 µL of 10x Nick translation buffer, with nuclease-free ddH2O added to a final volume of 45 µL. After vortexing and brief centrifugation, 5.0 µL of Nick translation enzyme was added. The mixtures were incubated at 15°C for 10 hours, heat-inactivated at 70°C for 10 minutes, and cooled to 4°C before transferring to RNase-free tubes. Following the addition of 1 µL of 0.5 M EDTA, 1 µL of linear acrylamide, 5 µL of 3M NaOAc (pH 5.2), and 125 µL of cold 100% ethanol, samples were incubated at -80°C for 2 hours. After centrifugation at 20,000 rcf for 60 minutes at 4°C, the pellet was washed with 1 mL of 75% ethanol, followed by centrifugation and drying at 37°C for 15 minutes. Finally, 50 µL of DEPC-treated water was added, and samples were vortexed at 200 rpm for 1 hour while stored at -20°C.

Cells were plated in a Poly-L-Lysine coated glass bottom 24-well plate (MatTek, P24GTOP-1.5-F). mESCs and OPCs were used at approximately 80% confluency. DEPC-treated 1x PBS was employed throughout. Freshly prepared 4% paraformaldehyde (PFA) in DEPC-treated PBS was added to each well for a 15-minute fixation. After washing with 1x PBS, cells were treated with warmed 200 mM VRC and subsequently washed three times. A 0.5% Triton X-100 buffer was added for 10 minutes on ice, followed by additional washes with PBS and saturation in 400 µL of 2x SSC for 2 hours at 37°C. Probes were prepared by mixing 100 ng of probe with 2 µL of competitor yeast tRNA, drying, and then dissolving in 10 µL of deionized formamide. Probes were denatured, and a hybridization buffer was created. The probe mixture was added to the cells, which were sealed and incubated for 16 hours at 37°C. Post-incubation, cells were washed with pre-warmed solutions in a series of washes, followed by incubation with DAPI for 5 minutes.

Lamin_B1 immunolabeling: 200 µl of blocking buffer (3% BSA, 10% NDS, 0.1% PBSTx) was added per well and incubated for 1 hr at RT. Lamin_B1 primary antibody (Abcam, Cat. ab16048) diluted (1:500) in 200 µl of blocking buffer was added to each well and incubated on a shaker for 48 hrs at 4 °C. Primary antibodies were rinsed off, and the cells were then washed with 0.1% PBSTx four times (10 min for each wash). The secondary antibody (Thermo Fisher, Cat. 31573) diluted (1:2,000) in 200 µl of the secondary antibody buffer (5% NDS in 0.1% PBSTx) was added to each well and incubated at RT on a slow rocker for 1.5 hrs. After the incubation period, secondary antibodies were rinsed off, and then the cells were washed with 0.1% PBSTx three times and washed once with 1X PBS.___Finally, 100 µL of ibidi mounting medium was added to each well, and plates were stored at 4°C until imaging.

### Gene Ontology, Gene Enrichment, and Gene Classification analysis

Gene ontology analyses were performed using ShinyGO (v0.77), a graphical enrichment tool [37] at Iowa University. GO biological process of gene sets was assessed by Fold Enrichment, setting the FDR cutoff to 0.05 and pathway sizes to a minimum 2 and a maximum of 2,000. The first 20 pathways are shown in the figures. The size of a data point (dot) represents the number of genes, and the color of a data point represents the -log10(FDR). Promoter enrichment in the ChIP data was assessed using ShinyGO (v0.77), considering upstream 300 bp as the promoter region. Statistical analyses were performed using Chi-squared tests comparing the characteristics of the gene sets with the remainder of the genes in the mouse genome. Data are presented as P value and FDR for each promoter. Gene class classification was performed using g:Profiler (version e102_eg49_p15_7a9b4d6) [38] and ShinyGO 0.77 webtools. Statistical analyses were performed using Chi-squared tests integrated in ShinyGO 0.77 webtool. Gene enrichment was performed using Enricher web-portal [39, 40]. We used the ENCODE histone modification 2015 database embedded in the Enricher web portal to assess the histone association for the input gene sets. The disease associations of the input gene sets were assessed using Clinvar data in the Enricher web portal.

### Protein Chip assay

Protein pull-down by RNA Pol II pSer2 was confirmed using the Agilent Protein 230 kit (Agilent, Cat. 5067-1517) according to the manufacturer’s protocol, and protein electroporation was performed using the 2100 Bioanalyzer. We used 8 µl elute per sample.

## Results

### The level of RNA Pol II pSer2 increases at the LB1 region in ESC-derived NPCs

To investigate whether actively transcribing genes are associated with the LB1 region, we first performed an immunofluorescence assay in mESCs and single clone derived (*in vitro*) NPCs using an RNA Pol II pSer2 antibody, a molecular marker of actively transcribed genes, and a LB1 antibody. We observed that, although the majority of the RNA Pol II pSer2 immunofluorescent signals were at the NI, a smaller fraction of RNA Pol II pSer2 signals overlapped with LB1 signals at the NP (Fig. 1A, 3X boxed areas, L and R, are enlarged 3X in the image panels), suggesting the possibility of actively transcribing genes at the LB1 region. When we quantitated the overlap of the RNA Pol II pSer2 and LB1 fluorescent signals using Manders’ Colocalization Coefficient (Fig. 1B, C) and Pearson’s Colocalization Coefficient (Fig. 1D), we found that a significant fraction of RNA Pol II pSer2 is colocalized with LB1 at the NP (Fig. 1b - ESC: Manders’ Colocalization coefficient 0.5 (SEM +/-0.07), NPC: Manders’ Colocalization coefficient 0.8 (SEM +/-0.02), Fig. 1c - ESC: Manders’ Colocalization coefficient 0.2 (SEM +/- 0.07), NPC: Manders’ Colocalization coefficient 0.4 (SEM +/-0.01), Fig. S1b - ESC: Pearson’s Colocalization coefficient 0.36 (SEM+/- 0.05), NPC: Pearson’s Colocalization coefficient 0.64 (SEM +/- 0.01). Interestingly, in both analyses we observed that the RNA Pol II pSer2 signals at the LB1 region was higher in NPCs than ESCs, indicating higher gene transcription at the NP in NPCs. In addition, in our immunofluorescent images (Fig. 1E), we also observed that large-dense chromatin territories (presumably LADs) were present at the NP (arrows in yellow boxes) which were lacking the RNA Pol II pSer2 signal, which was mostly present at less dense chromatin territories (arrows in red boxes).

**Fig. 1.**
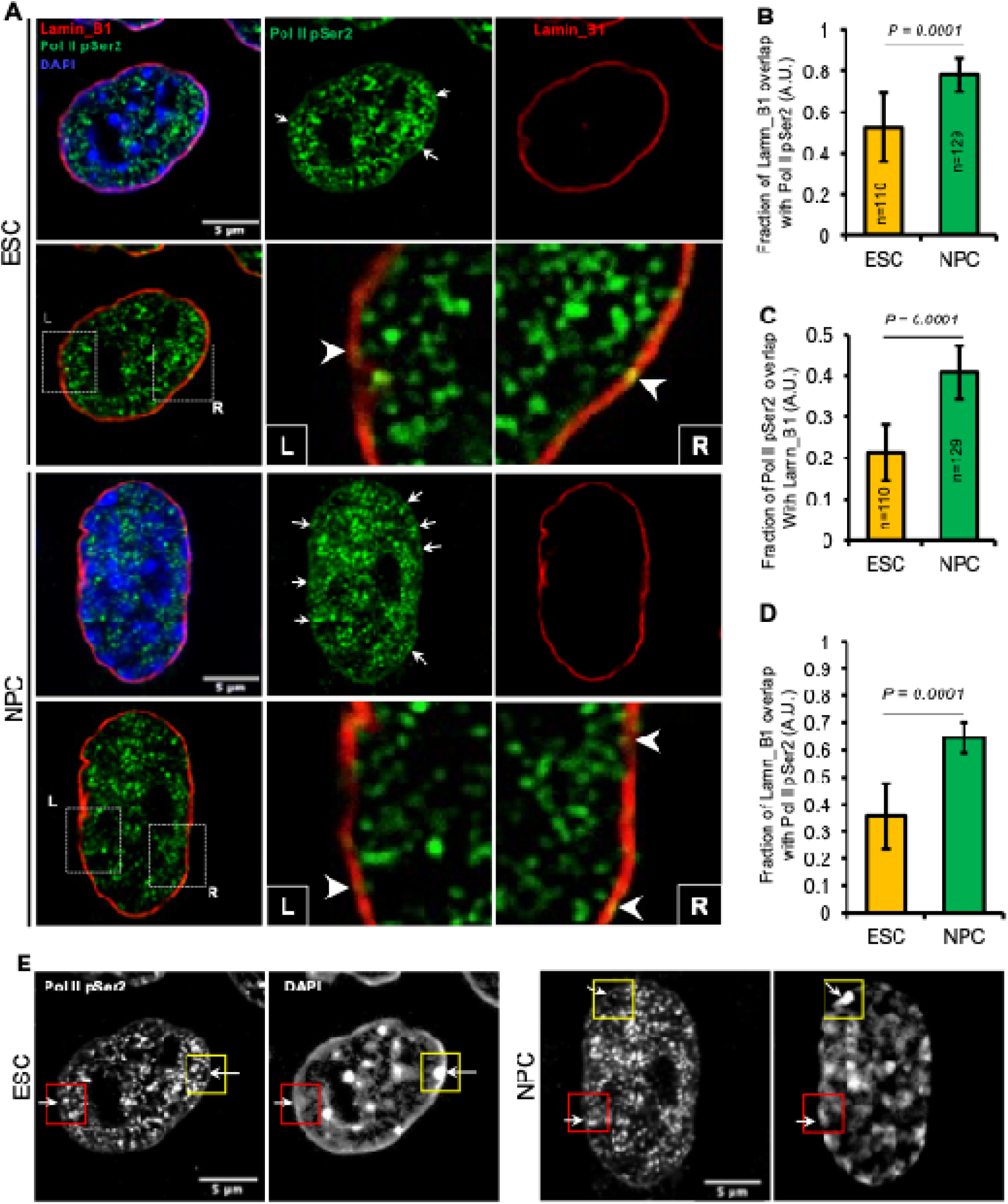
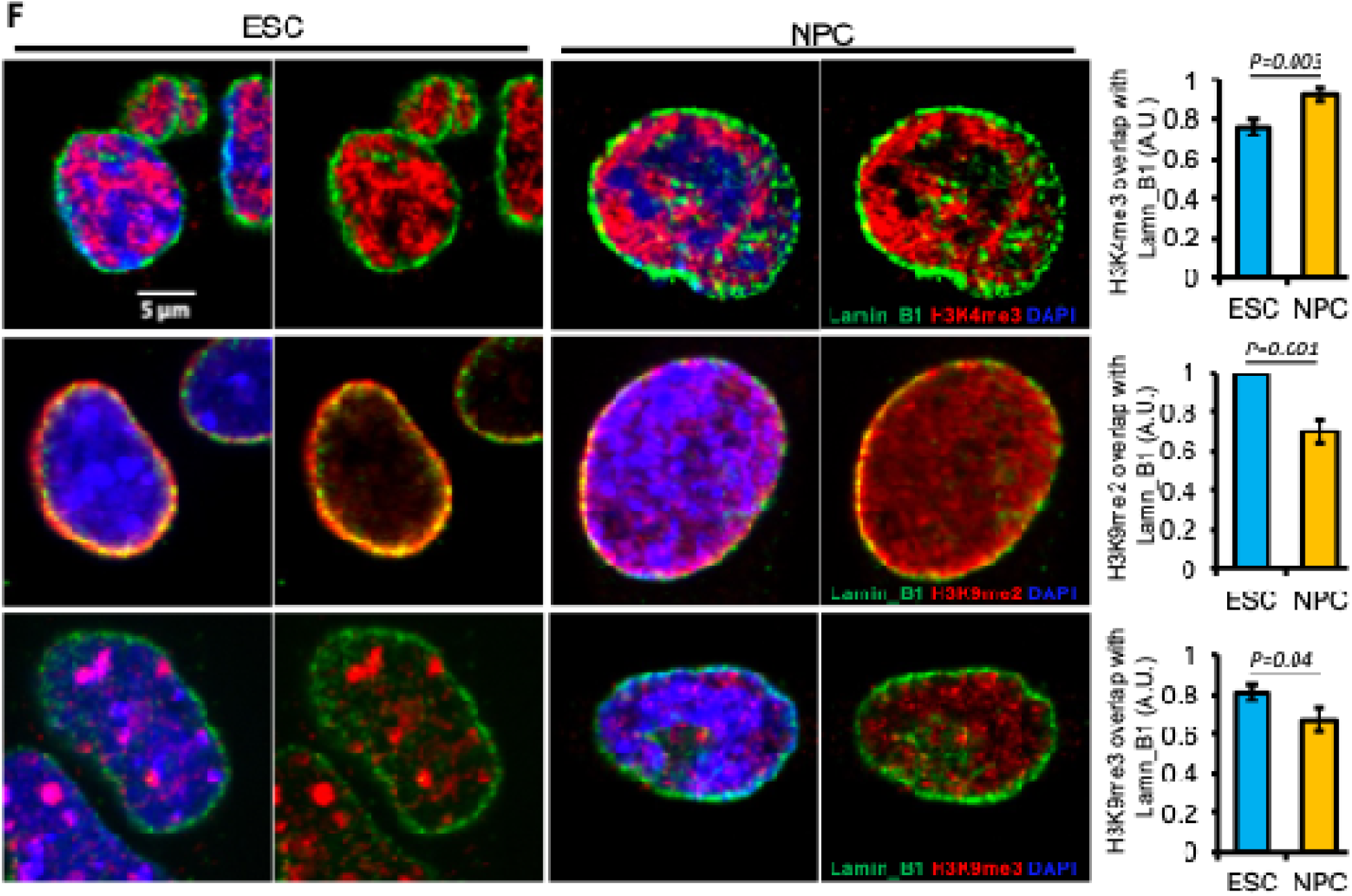
Presence of LB1 associated RNA Pol II pSer2 increases at the NP in mESCs-derived NPCs. **A** Representative confocal microscopy images showing LB1, RNA Pol II pSer2 and DAPI staining in mESCs (upper panel) and NPCs (lower panel) (Scale bar – 5 μm). Images show LB1 localization along the nuclear periphery and RNA Pol II pSer2 sporadic distribution in the cells, mainly in the nuclear interior (NI) and some are overlapped with and adjacent to LB1. Single channels (RNA Pol II pSer2 and LB1) fluorescent images showing the localization of each protein. White arrows showing some of the RNA Pol II pSer2 (green) signals at the NP of ESCs and NPCs. Visually, more green fluorescent signals are present at the NPCs NP than in ESCs NP. The left (L) and right (R) boxed images in each panel are enlarged 2.5X. Arrowheads in the last two images in both rows show the LB1 and RNA Pol II pSer2 overlaps. **B** Bar-graph showing the Manders’ Colocalization Coefficient of LB1 fluorescent signals overlapping with RNA Pol II pSer2 fluorescent signals calculated using ESC and NPC images. Colocalization of LB1 and RNA Pol II pSer2 fluorescent signals are significantly different (p<0.0001) between ESCs and NPCs. **C** Bar-graph showing the Manders’ Colocalization Coefficient of RNA Pol II pSer2 fluorescent signals overlapping with LB1 fluorescent signals calculated using ESC and NPC images. Colocalization of RNA Pol II pSer2 and LB1 fluorescent signals are significantly different (p<0.0001) between EScs and NPCs. (ESC: n=110, NPC: n=129, A.U. – Arbitrary Units). **D** Bar-graph showing the LB1 and Pol II pSer2 colocalization analysis results using Pearson Colocalisation Coefficient. Colocalization of RNA Pol II pSer2 and LB1 fluorescent signals are significantly different (p<0.0001) between ESCs and NPCs. (ESC: n=110, NPC: n=129, A.U. – Arbitrary Units). **E** Gray channel images showing Pol II pSer2 and DAPI (chromatin) localization in ESCs and NPCs. The arrows in the yellow squares show chromatin areas lacking RNA Pol II pSer2 in the NP. Arrows in red squares show chromatin areas with RNA Pol II pSer2. (Scale bar – 5 μm). **F** Representative microscopic images showing the Lamin_B1, H3K4me3 (top panel), H3K9me2 (middle panel) and H3K9me3 (lower panel) labeling in mESCs and NPCs (Scale bar – 5 μm). Images were quantified for Lamin_B1 and each histone mark to assess their colocalization. The bar graph corresponding to each raw image shows the colocalization of both signals. Manders’ Colocalization Coefficient method was used for analysis.

H3K4me3 is known to mark transcriptionally active genes [41] and H3K9me3 marks the transcriptionally repressed genes [42]. Specifically, H3K9me2 is known to mark the transcriptionally inactive genes at the nuclear periphery [43]. Therefore, to further validate our above results, we used these three histone marks to assess their colocalization with LB1 by immunofluorescence and image analysis (Fig. 1F). Assessing the colocalization of LB1 and each histone mark using Manders’ Colocalization coefficient, we found that H3K4m3 signals were associated with LB1 and significantly increased upon ESC differentiation to NPCs (p=0.001) suggesting an increase in transcriptionally active genes at the nuclear periphery in NPCs (bar graphs at the end of the top image panel). In contrast, compared to ESCs, H3K9me2 and H3K9me3 signals associated with LB1 at the nuclear periphery in NPCs was significantly decreased (P=0.001 and P=0.04, respectively), confirming the decrease in transcriptionally active genes at the LB1 compartment of NPCs (bar graphs at the end of the middle and lower image panels).

### Sequential ChIP-seq analysis revealed that a sub-set of actively transcribed genes are present at the LB1 nuclear region

RNA Pol II pSer2 marks actively transcribing gene bodies [44]. Therefore, based on our immunofluorescence observation, we sought to investigate the RNA Pol II pSer2-associated genes at the LB1 compartment in a high-throughput manner. To do so, we utilized a sequential ChIP assay, first to isolate all the genes associated with the LB1 compartment and then to specifically pull down the genes that interact with RNA Pol II pSer2; therefore, actively transcribing genes in the LB1 compartment (Fig. 2A). In the first step of the sequential ChIP assay, we pull down ∼83% of the total LB1 protein in the input lysate using LB1 antibody (Fig. 2B, C) from mESCs. We then used the pulled-down LB1-chromatin fraction as the input to pull down the genes bound by RNA Pol II pSer2. Next, to identify the genes that were enriched in the RNA Pol II pSer2 pull-down chromatin, we constructed DNA libraries using the Illumina ChIP-seq kit and sequenced the DNA using the Illumina HiSeq sequencing platform. We used 150 paired-end sequencing to increase the alignment accuracy and minimize the probability of aligning into multiple genomic loci when aligning to the reference genome.

**Fig. 2.**
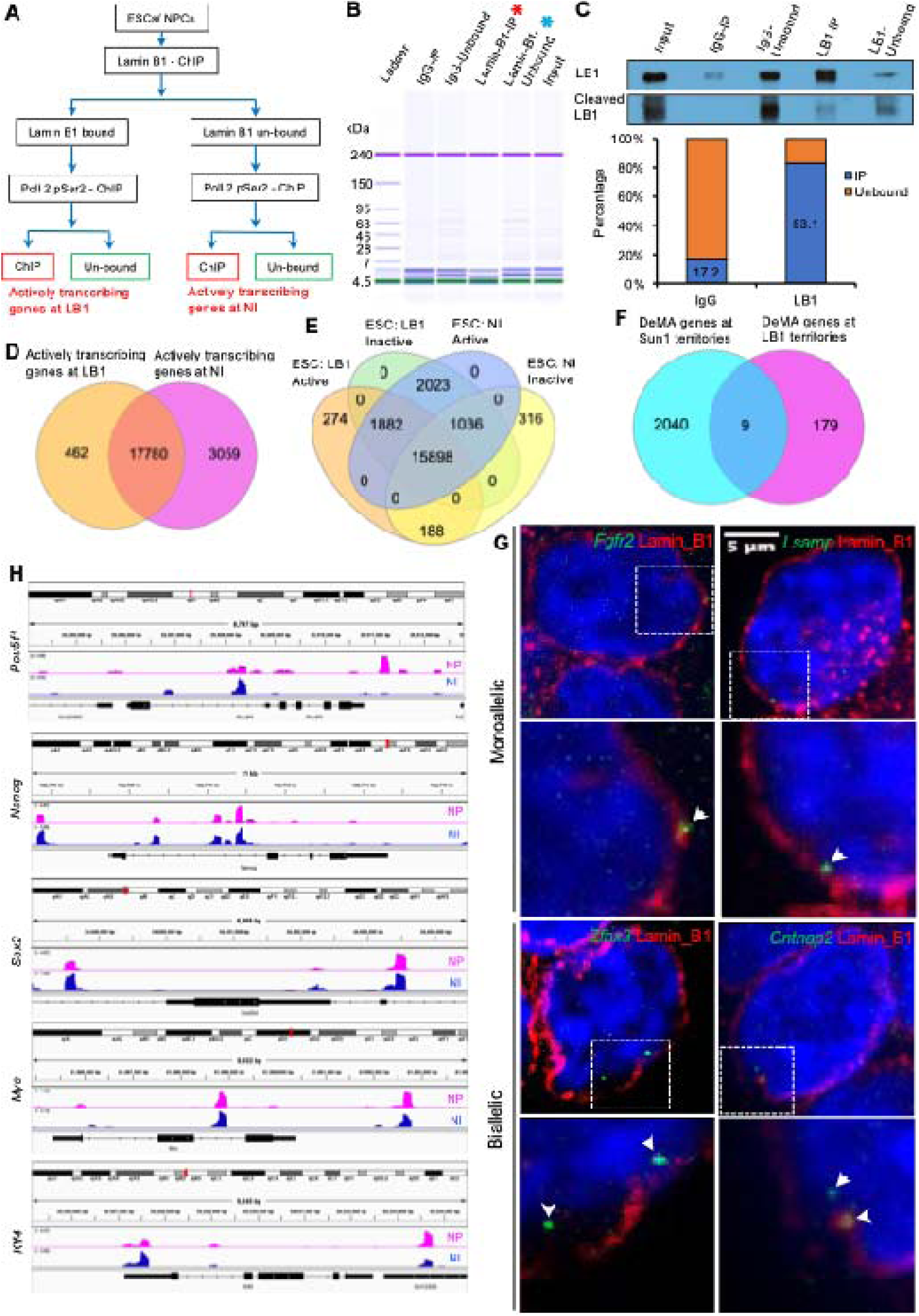
Sequential ChIP assay revealed genes that are actively transcribed at the LB1 nuclear region. **A** Schematic presentation of sequential ChIP workflow. In the first step of the pull-down assay, LB1 bound and unbound fractions were separated and used as the input material for the second step using RNA Pol II pSer2 antibody. DNA was extracted from the chromatin fraction pull-down by pSer2 (red rectangles in the diagram) and the pSer2 unbound (green rectangles in the diagram) chromatin. (LB1: Lamin_B1 compartment, NI: nuclear interior). **B** Protein content in different fractions in the first step (pull-down by LB1) of the sequential chip assay in ESCs. Only the LB1 bound (red star) and LB1 unbound (blue star) protein-chromatin fractions were used for the second step of the pull-down assay. The protein contents in the LB1 pull-down fractions are very low compared to the LB1 unbound and IgG control fractions. **C** Immunoblot showing the specificity of the LB1 pull-down in the first step of the sequential ChIP assay. LB1 unbound fraction shows that LB1 protein was not completely pulled down and remains in the LB1 unbound fraction. The cleaved LB1 protein fraction is higher in the LB1 unbound fraction compared to the LB1 and IgG-bound fractions. The bar graph shows the percentages of the quantitated LB1 signal intensities of the bands in bound and unbound fractions. Total intensities were calculated by summing the signal intensities of the LB1 band and the cleaved LB1 band in each lane. **D** Venn diagram showing the actively transcribing genes in LB1 compartment and NI compartment of mESCs assessed by ChIP-seq analysis. 18,242 and 20,839 actively transcribing genes were detected in LB1 and NI compartments, respectively, and 17,780 genes are common to both gene categories. 462 genes are exclusively expressed in the LB1 compartment, while 3,059 genes are exclusively expressed at the NI. **E** Venn diagram demonstrating the monoallelically (188) and biallelically (274) expressed genes in the LB1 compartment in mESCs. **F** Venn diagram showing the intersection of DeMA (deterministic monoallelic genes) genes at the Sun1 territories at the inner nuclear membrane and the LB1 region. The results show that a very low percentage of genes are common to both compartments. **G** Nascent RNA-FISH images showing genes are actively transcribed at the LB1 region in ESCs. Four genes were selected from the sequential ChIP-seq data based on higher read counts. *Fgfr2* and *Lsamp* genes are known to be expressed monoallelically in ESCs [32] and bi-allelically expressed *Klf3* and *Et14* genes; both alleles are in the same x-y image plane. Images in the lower panels are the 2.5X magnified boxed area in each image of the top image panels. (Scale bar – 5 μm). White arrowheads show the nascent RNA-FISH signals. Transcripts are labeled in red and LB1 is labeled in green (overlap is yellow). Nuclei are labeled with DAPI. **H** ChIP-seq peak tracks for *Pou5f1*, *Nanog*, *Sox2*, *Myc* and *Klf4* genes illustrated in integrative Genomic Viewer. Data summarized in each peak as the mean of all the pixels overlapped into a single genomic location. Magenta-colored tracks show the transcribing genes/alleles in the LB1 compartment, and the blue-colored tracks show the genes/alleles transcribed at the NI. The *Pou5f1* gene is mainly transcribed in the NI, and *Nanog* is transcribed either in the LB1 compartment or the NI compartment. Scales for both tracks were set to auto scale.

In our ChIP-seq analysis, we found that a total of 21,301 genes were expressed (RNA Pol II pSer2 bound) in mESCs, and from that, 2.7% (462) of genes were actively transcribing at the LB1 region (Fig. 2D). We found that 17,780 genes are common to both the LB1 compartment and the NI in mESCs. We speculate that this may be due to incomplete LB1 pull-down in the first step of the sequential ChIP assay (Fig. 2C) or each of the two alleles of a gene being expressed in one of the two nuclear regions, LB1 or NI. Alternatively, some genes may not have preferential compartmentalization (LB1 or NI) in the nucleus when transcriptionally active. However, we cannot provide a definite explanation for those genes common to the LB1 and the NI regions since our data is from bulk RNA-seq analysis. Therefore, we did not include genes common to both the LB1 and NI compartments for the downstream analysis to avoid misinterpretation of our ChIP-seq data. To assess the genes that are exclusively expressed in the LB1 region, we intersected the active (LB1 and RNA Pol II pSer2 associated) and inactive (LB1 associated but pSer2 negative) gene sets at the LB1 region, and the active (NI and RNA Pol II pSer2 associated) and inactive (NI associated but RNA Pol II pSer2 negative) gene sets at the NI from mESCs. We found that both alleles of 274 genes were expressed at the LB1 region, while for an additional 188 genes, only a single allele was expressed at the LB1 region in mESCs (Fig. 2E).

We have previously shown that many monoallelic genes are preferentially expressed at Sun1-enriched territories at the NP [33]. Therefore, we asked whether the monoallelic genes expressed at the Sun1 territories and the LB1 compartment-associated monoallelic genes in mESCs were common. We found that only ∼5% of the transcriptionally active genes from the LB1 region and the Sun1 territories overlapped (Fig. 2F), suggesting that the LB1 region and Sun1 territories may differentially scaffold the monoallelic genes at the NP. However, given that we could only pull down ∼83% of the LB1 protein at the first step of the sequential ChIP assay in mESCs, some of the active monoallelic genes associated with LB1 may not have been captured. Further, to validate the presence of actively transcribing genes at the LB1 region, we performed nascent RNA-FISH for four highly expressed (high read counts) gene candidates. Confirming our finding from the sequential ChIP-seq assay, we observed the tested genes are actively transcribed at the LB1 region (Fig. 2G), and the monoallelic expression of the *Fgfr2* and *Lsamp* genes agrees with our previously published data [32].

Genes may be organized in different nuclear regions depending upon the state of the cell [45, 46]. Therefore, we asked whether any known genes involved in pluripotency (*Pou5f1, Nanog, Sox2, Klf4*, and *Myc*) were actively transcribed at the LB1 compartment in mESCs. However, we observed that only the *Pou5f1* gene’s alleles are predominantly expressed in the NI, but the rest of the genes’ alleles could be either at the NP or NI (Fig. 2H). Further, to understand whether the genes localized at the LB1 region play a significant role in mESC specific functions, we searched for the gene ontologies associated with the actively transcribed genes at the LB1 compartment. However, we did not observe any significant gene ontology enrichment specific to stem cells. Instead, these genes consisted of housekeeping genes.

### Actively transcribed genes proportionally increase their occupancy at LB1 in NPCs primed toward the OPC lineage

It has been previously shown by RNA-seq analysis that when mESCs differentiate into NPCs *in vitro*, the total number of expressed genes declines by ∼50% [47]. Therefore, to test whether we could utilize our sequential ChIP assay to recapitulate the reported gene expression changes in NPCs, we pulled down the RNA Pol II pSer2-associated chromatin and assessed the total number of RNA Pol II pSer2-associated genes in mESCs versus NPCs (Fig. 3A-C). Using our assay, we observed that compared to mESCs, the total number of genes expressed in NPCs had declined by nearly 28% (Fig. 3D – Blue columns), showing the same gene expression trend as previously reported [47]. Further, it has been shown that when cells differentiate, chromatin organization in the nucleus changes [48-52]. Moreover, a study using the LB1 Dam-ID method has shown that the number of heterochromatic genes increases at LADs in differentiated cells vs undifferentiated naïve cells [53]. However, in our immunofluorescence assay, we observed that the percentage of the RNA Pol II pSer2 signals at the LB1 region was increased in NPCs compared to mESCs (Fig. 1B-D). When we assessed our ChIP-seq data for gene enrichment in the LB1 region, aligned with the immunofluorescence observation, we found that the number of actively transcribing genes at the LB1 region was also increased by ∼20 fold in NPCs (Fig. 3D- orange color columns, E). Further, using the nascent RNA-FISH assay for four genes, we also validated that these genes were actively transcribed at the LB1 region (Fig 3G).

**Fig. 3.**
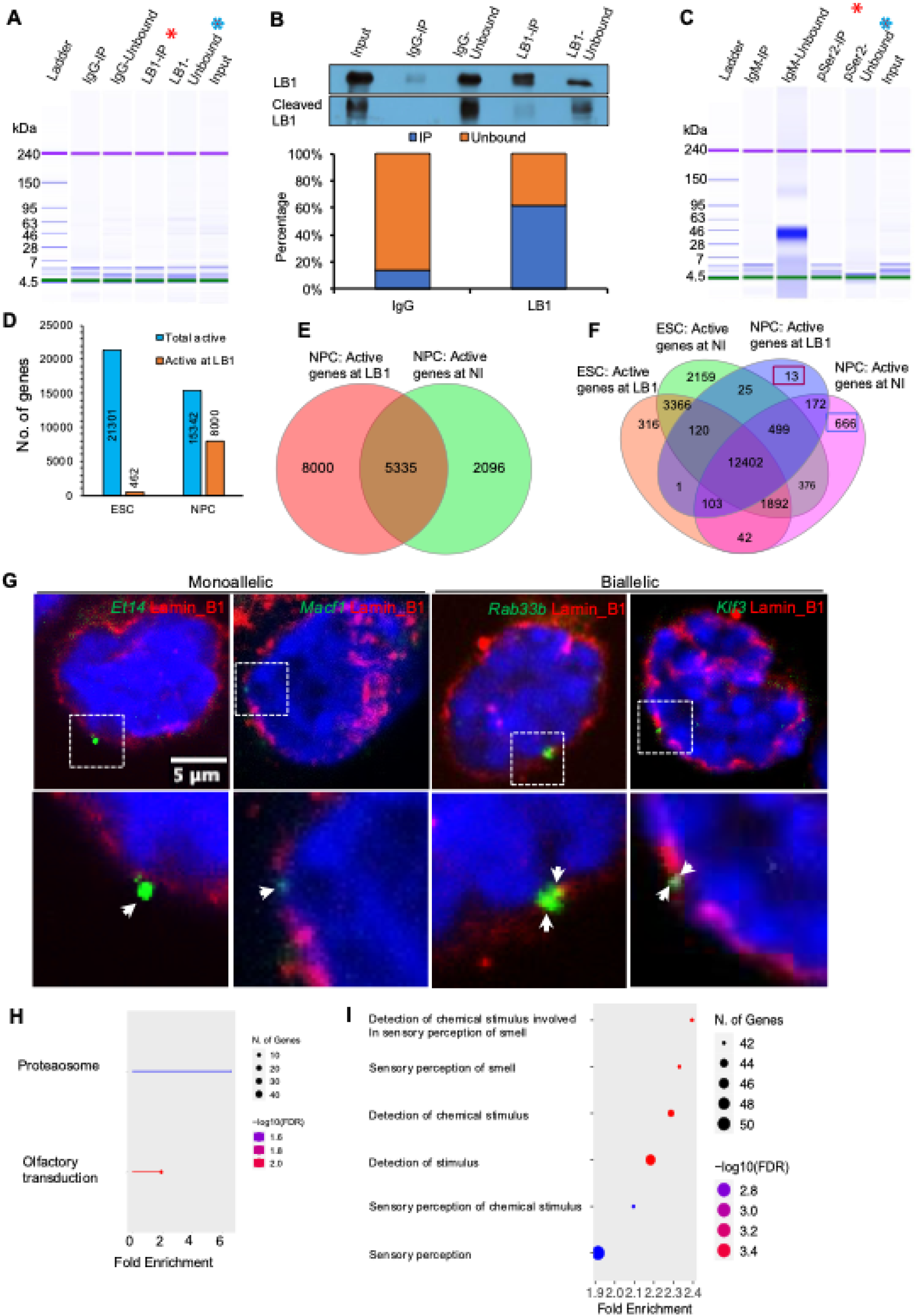
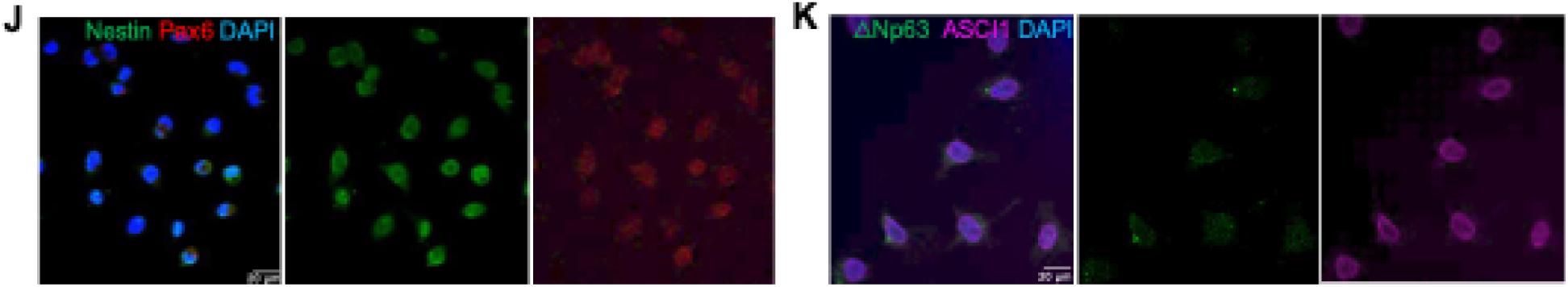
Actively transcribed genes proportionally increase their occupancy at LB1 in mESC-derived OPCs. **A** The protein chip shows the relative protein fractions pulled down by LB1 in the first step of the sequential ChIP assay using NPCs. Only the LB1 bound (red star) and LB1 unbound (blue star) protein-chromatin fractions were used for the second step of the pull-down. **B** The representative immunoblot is showing the LB1 pull-down efficiency in NPCs. Compared to the unbound fractions of IgG and LB1, the signals relevant to cleaved LB1 are lower in IgG and LB1 fractions, similar to the observation made in mESCs. The histogram shows the percentages of signal intensities of IgG and LB1 pull-down bands. Percentages were calculated over the total LB1 (in each IgG and LB1 fraction), and the totals of uncleaved and cleaved signals were summed to have the total LB1 signals. In NPCs, ∼62% of the total LB1 protein is pulled down. **C** Protein contents in different fractions in the second step of the sequential ChIP assay in NPCs. **D** The bar chart shows the total number of actively transcribing genes (Blue-colored) and the actively transcribing genes at the LB1 compartment and NI in mESCs and NPCs assessed by ChIP-seq assay. The total number of actively transcribing genes declines by ∼28% in NPCs compared to ESCs. The actively transcribed genes at the LB1 compartment in NPCs have increased by ∼20% from the total actively transcribed genes in ESCs. **E** Venn diagram showing the intersects between actively transcribed genes at the LB1 compartment and NI in NPCs. The number of genes expressed from both compartments is less than in ESCs. **F** Venn diagram shows the intersects of actively transcribed genes compartmentalized in ESCs and NPCs. Genes that are uniquely expressed at the LB1 (red rectangle) and NI (blue rectangle) are indicated in the diagram. **G** Nascent RNA-FISH images showing validation that genes are actively transcribed at the LB1 region in NPCs. Four genes were selected from the sequential ChiP-seq data based on higher read counts. Images in the lower panels are the 2.5X magnified dotted-lined area in each image of the top image panel. (Scale bar – 5 μm). White arrows show the nascent RNA-FISH signals. The four genes are bi-allelically expressed, and both alleles are in the same x-y image plane. **H** KEGG analysis for the actively transcribing newly established genes at the LB1 region in NPCs. Genes are only enriched for olfactory transduction and proteaosome pathways. The size of the dots indicates the number of genes in each category, and the colour of each dot indicates the -Log10 of FDR. **I** Gene Ontology analysis of the genes exclusively actively transcribed at NI in NPCs. The gene set is exclusively enriched in the stimuli-sensing gene ontology. The size of the dots indicates the number of genes in each GO category, and the color of each dot indicates the -Log10 of FDR. **J** IF images showing Nestin and Pax6 expression in the mESC-derived NPCs. The NPCs are positive for both Nestin and Pax6. Scale bar: 20 µm. **K** IF images showing the expression of △Np63 and Ascl1in NPCs. The NPCs are only positive for Ascl1; therefore, they are Globose basal (olfactory progenitor in early developing embryo’s olfactory epithelium) cells. (Scale bar - 20 µm).

When mESCs differentiate into NPCs, we observed 8,000 genes (423 genes in mESCs) established their transcription at the LB1 region. Therefore, to investigate whether the increased gene activity at the LB1 compartment in NPCs is linked to the specific onset of lineage commitment of these cells, we assessed the gene ontologies of the newly established actively transcribing genes at the LB1 region (13 genes) and the genes uniquely expressed (851 genes) in NPCs (Fig. 3F). For the newly established actively transcribing genes at the LB1 region in NPCs (13 genes), we did not observe any neural-related or cell lineage-specific gene enrichments; instead, we found that those genes exhibit “housekeeping” molecular functions. However, for the total newly established actively transcribing genes in NPCs (851 genes), we found that KEGG pathways are only enriched for olfactory transduction (fold enrichment: 1.9, gene enrichment FDR: 7.2E-03) and proteasome (fold enrichment: 6.5, gene enrichment FDR: 3.4E-02) pathways (Fig. 3H). In addition, testing the genes actively transcribing only at NI regions (666 genes), we found that those genes were enriched in the sensory perception and stimulus detection pathways (Fig. 3I).

Although it is generally accepted in most mammalian cells that LB1 regions predominantly harbor heterochromatin, chromatin organization in the nuclei of olfactory neurons has been shown to have an inverse heterochromatin and euchromatin organization [22]. Therefore, based on the gene ontologies of the newly established actively transcribing genes in clones derived from NPC cultures, we speculated that some cells in NPC cultures might represent primed precursor olfactory neuron cells of which euchromatin and heterochromatin arrangements were predetermined. Although Pax6 and Nestin are known neural progenitor markers, it has also been shown that Nestin is expressed in radial glia-like progenitors that differentiate to zonally restricted olfactory and vomeronasal neurons [54] and that Pax6 is critical in neuronal incorporation into the adult olfactory bulb [55]. Interestingly, the cells in our NPC cultures are positive for both markers (Fig. 3J). Therefore, we assessed the OPC-specific signature in our NPC cultures using the olfactory progenitor cell markers Ascl1 (Globose basal cells-GBCs) and △Np63 (Horizontal basal cells-HBCs). GBCs originate in early development, and the HBCs appear in the olfactory epithelium at a later developmental stage [56]. In our immunofluorescence assay for Ascl1 and △Np63, we observed that the OPCs are predominantly positive for Ascl1 but with varying levels of labeling in a few OPCs, confirming that our *in vitro*-derived NPCs mimic the cellular identity of olfactory progenitors of the embryonic placode [57] (Fig. 3K).

### Actively transcribing genes at the LB1 region exhibit unique molecular characteristics

RNA Pol II is known to transcribe protein-coding and non-coding RNA genes [58]. Therefore, to investigate whether there are preferentially transcribing gene classes at the LB1 region, we next assessed the enrichment of gene classes among the LB1 associated actively transcribing genes (mESCs - 423 genes and NPCs - 8000 genes). We found that the input gene sets represent genes from various gene classes, and the enrichment of some classes was seemingly increased when mESCs differentiated into OPCs (Fig. 4A). Furthermore, those LB1-associated actively transcribing genes were spread across the genome (Fig. 4B) and we found that more genes were accumulated around those "preset loci" in mESCs when differentiated to OPCs (Fig. 4B – e.g., dotted blue rectangles on Chr2 and Chr19).

**Fig. 4.**
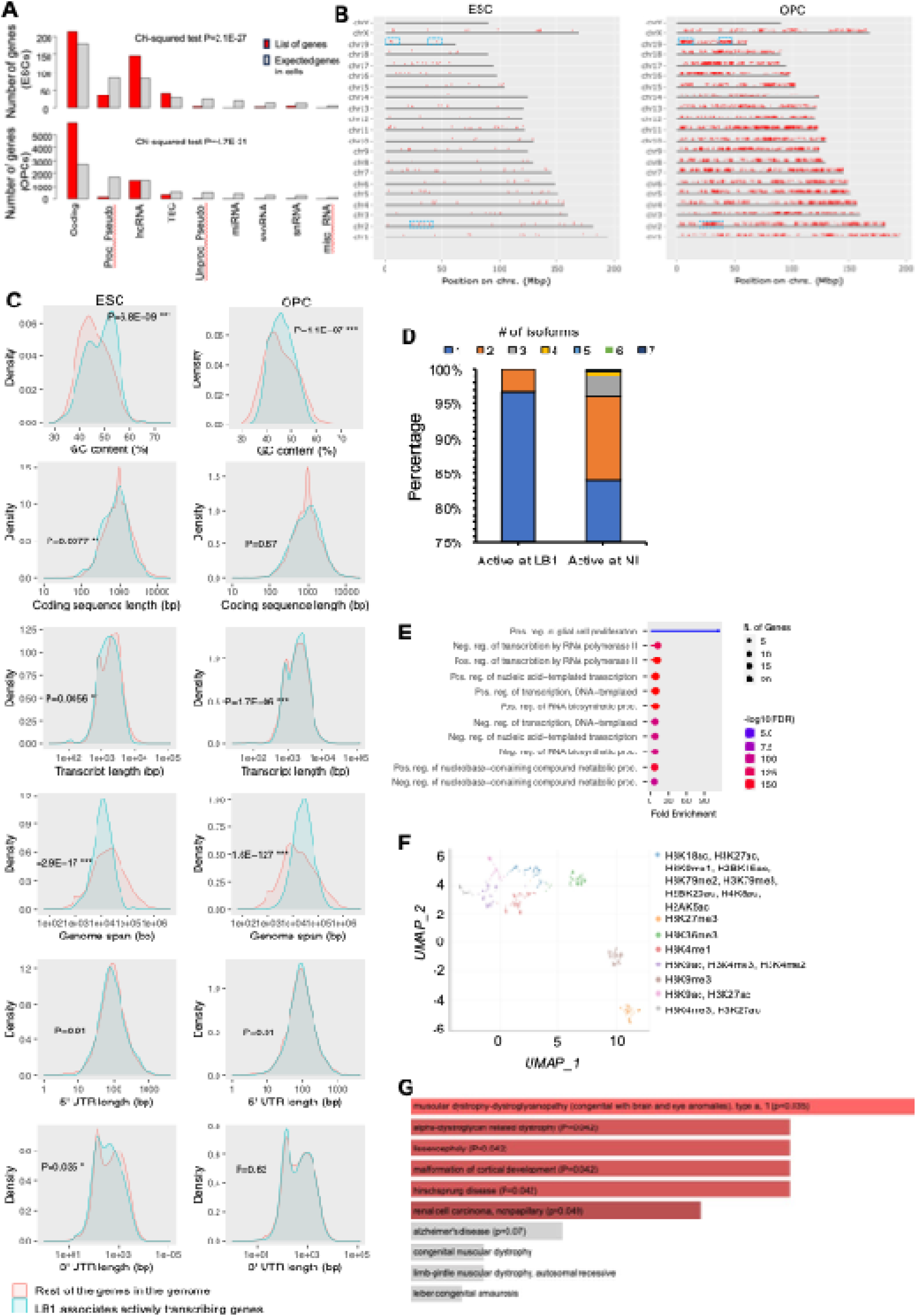
Actively transcribed genes at the LB1 compartment exhibit unique molecular characteristics. **A** The bar plots summarize the classification of the actively transcribed genes in the LB1 region in ESCs and OPCs. Statistical analysis was performed using the Chi-squared test integrated in ShinyGO 0.77 software. **B** Illustration of RNA Pol II pSer2 pull-down gene distribution through the genome. Compared to ESCs, there is an accumulation of actively transcribing genes at the LB1 compartment in OPCs. Also, in OPCs, more genes accumulate around the gene loci, which were already established at the LB1 region in ESCs (e.g., Chr2 and Chr19 – blue rectangles). **C** Density plot showing molecular characteristics of the genes actively transcribing at LB1. Guanin and cytosine is significantly enriched at actively transcribing genes at the LB1 region and the transcript length and genome span significantly differ from the rest of the genes in the genome compared to the rest of the genes in the genome in both ESCs and OPCs. **D** Histogram shows the number of isoforms per gene in LB1-associated versus NI-associated genes. The data indicate that the LB1-associated genes exclusively have either one or two known isoforms, while actively transcribing genes at the NI could have more than two isoforms. **E** Gene Ontologies of the transcription factors known to bind to the promoters of the actively transcribing genes at the LB1 region. GO terms exhibit that those TFs are associated with RNA Pol II. The size of the dots shows the number of genes, and the color of the dots shows the -Log10 of the FDR of each GO term. **F** In silico analysis of histone signature association with the actively transcribing genes at the LB1 compartment. The dots in the UMAP plot indicate the known histone signatures associated with the assessed genes. “Blue” dots indicate the known histone signature of active gene bodies, as well as active enhancers (H3K27ac, H2BK20ac, HeK79me2), active promoters (H4K8ac, H2AK5ac, H3K9ac), and TSS of the transcribing genes (H3K4me3, H3K4me2). **G** Bar plot showing the gene set association in Clinvar data. The X-axis represents the -log10 of the p value, and the actual p values are indicated in each column. The red color shows the enriched terms for the input gene set from mESCs and OPCs.

Furthermore, to understand the molecular features of the actively transcribing genes at the LB1 region, we next investigated the molecular characteristics of those genes, focusing on the coding sequence length, transcript length, genome span, 5’ UTR length, 3’ UTR length, and guanine and cytosine (GC) content (Fig. 4C). Compared to all other genes in the genome (tested by Chi-square and Student’s t-tests), we found that the genome span, transcript length and the GC content of the actively transcribing genes (mESCs: 462 and OPCs: 8000) at the LB1 region in mESCs and OPCs were significantly higher (ESC: p=6.8E-09 and OPCs: p=1.1E-07) compared to the rest of the genes in the genome. Moreover, we found that actively transcribing genes at the LB1 region transcribe a single or two isoforms, while the actively transcribing genes at the NI were generally found to transcribe more than two isoforms (Fig. 4D).

It has also been shown that the nuclear envelope could contain resting transcription factors (TFs) [59], and TFs interacting with Nups at the nuclear pore complex could play a role in gene localization, transcription repression, transcription poising, and transcription activation (reviewed in [60]). Therefore, taking this as a lead, we assessed the molecular features of the promoters of LB1-associated genes and the known TFs bound to those promoters. We found that the genes exclusively transcribed at the LB1 region were enriched with G-rich promoters (Table 1). Further, promoter motifs in the rest of the mouse genome similar to the motif associated with our gene sets are known to be regulated mainly by zinc finger TF family proteins (Table 1), and these TFs are known to regulate RNA Pol II-dependent gene transcription of both protein-coding and non-coding genes (Fig. 4E).

**Table 1:**
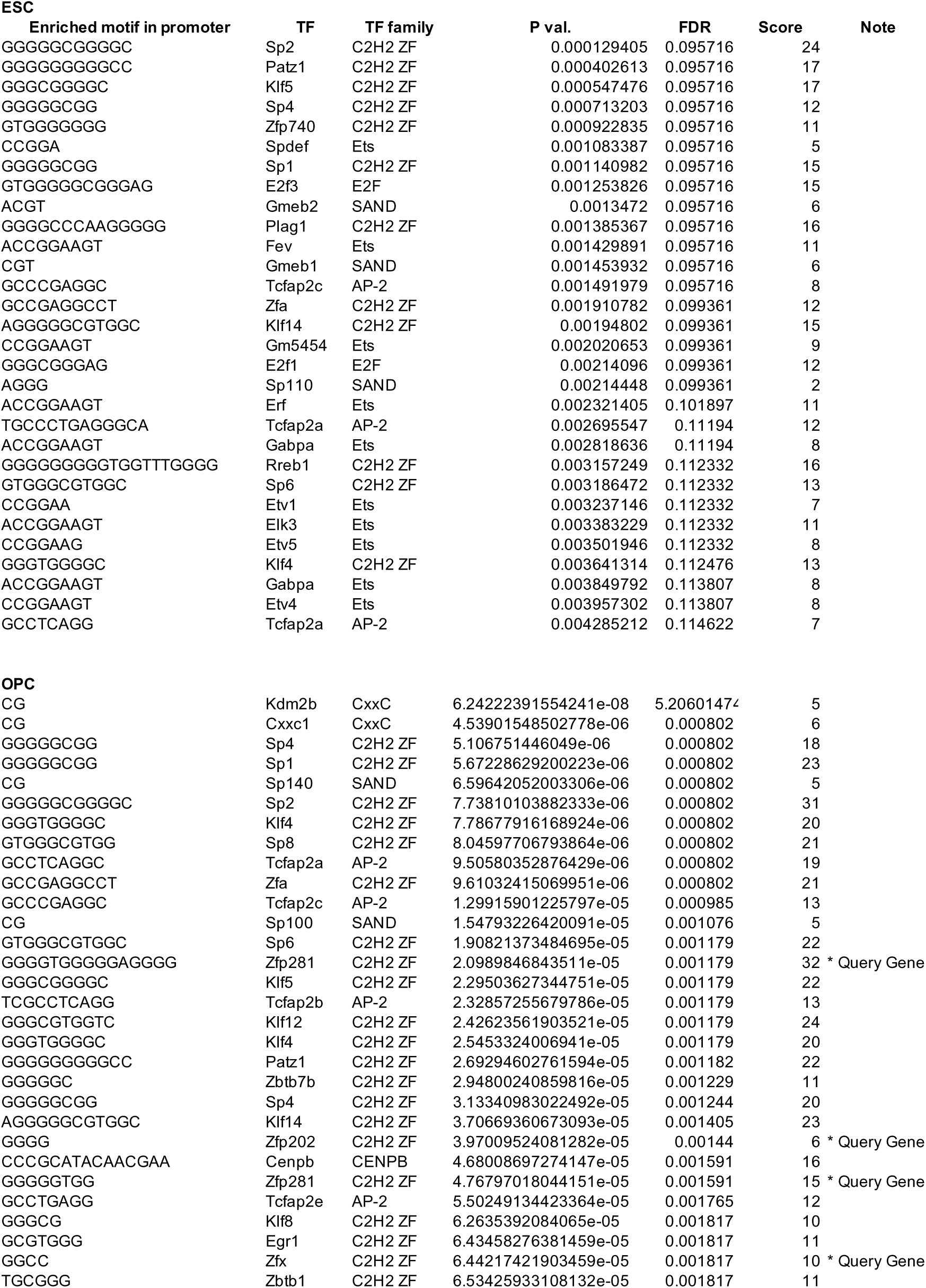
Enriched motifs in promoters and the promoter interacting TFs of the genes actively transcribing at the Lamin B1 region.

Because the actively transcribing genes at the NP found in this study were associated with LB1, we investigated the known histone signatures enriched for those genes using the ENCODE database embedded in Enricher - a data analysis web tool [61-63]. We found that the genes we tested exhibit associations with known histone signatures (H3K36me3, H2BK15ac, H3K79me2, H3K79me3, H4K8ac, H3K4me3), which mark actively transcribing gene bodies (Fig. 4F).

## Discussion

Although LADs are known to predominantly harbor heterochromatin, growing evidence suggests that LADs are not completely heterochromatic territories but do harbor active genes [15, 19, 64, 65]. In support of this idea, Leemans and colleagues [66] have shown that repressed promoters in LADs can become activated, however the mechanism is not yet known. In the present study, immunofluorescence data in mESCs and OPCs strongly suggests that actively transcribed genes (marked by RNA Pol II pSer2) are present at the NP associated with LB1. To validate this initial observation we utilized a sequential ChIP-seq assay to investigate the enriched active genes associated with the LB1 region and we further validated several gene candidates from the ChIP-seq assay by combining nascent RNA-FISH and LB1 immunofluorescence in mESCs and OPCs. Using three distinct experimental approaches, we demonstrated that LB1 harbors actively transcribing genes in mESCs and OPCs. In line with our finding of transcriptionally active genes at the LB1 compartment, the increase in transcriptionally active genes at the OPCs’ LB1 compartment indicates that, because the mature olfactory cells are programmed to establish a heterochromatic NI compartment, the establishment of housekeeping genes at the Lamin_B1 compartment is a unique chromatin organization phenomenon specific to olfactory cell lineage differentiation. It has been shown that upon cell differentiation into more specialized cells, more genes tend to be repositioned at the inner NE and gain heterochromatic status [53]. However, in line with the fact that euchromatin and heterochromatin organization is inverted in olfactory cell nuclei [21], our findings support the emerging idea that the LB1 compartment is not solely heterochromatic but can also accommodate transcriptionally active genes based on the cellular context. Not only in olfactory receptor cells, but also in other cell types, such as B-cells [67] and rod cells [68], chromatin organization similar to olfactory cells has also been reported. Therefore, because different cell types can have different chromatin architectures, the organization of transcriptionally active and silent genes needs to be assessed within the context of a specific cell type being studied and at the individual gene level.

The RNA Pol II pSer2 bound, therefore actively transcribing, genes we found at the LB1 region contain unique molecular characteristics, for example, high GC enrichment, higher genomic span, and long transcript length. Genes that transcribe longer transcripts have been shown to gain fewer SNPs through evolution than shorter genes [69-73]. Therefore, since the LB1-associated genes we identified in our study are housekeeping genes that share similar molecular functions across all cell types, it is reasonable to argue that these genes may be less prone to change during gene evolution. Although longer genes are associated with embryonic development [74] and neuronal development [75], we found no cell type specificity relating to the LB1-associated actively transcribing genes.

Furthermore, the relative GC enrichment of a gene is positively correlated with the transition of double-stranded DNA from its B form to Z form, increasing the bendability and reducing the thermostability of the DNA [76]. In addition, increased GC content is consistent with an open chromatin structure supporting stable and continuous transcription, which would be beneficial for housekeeping genes. Furthermore, open chromatin provides the advantage of minimizing the energy cost to the cell by reducing frequent histone modifications, which require energy usage. Therefore, the evolutionary advantage of high GC content in these genes may be used as a mechanism that could result in continuous open chromatin with minimal involvement of histone modification changes, a hypothesis worth examining in detail in future studies.

By compiling the LB1-associated genes with ENCODE histone data, we observed that histone signatures known to mark actively transcribing gene bodies are associated with genes we pulled down with RNA Pol II pSer2 from the LB1 region. Along with H3K27ac and H2BK20ac, we observed that H3K79me2, an intergenic enhancer signature, is also associated with the LB1 region-associated genes (Fig. 4F). H3K79me2 is known to facilitate chromatin accessibility for transcription factors [77], thus further validating the transcriptional activity of the LB1 region-associated genes we found. However, as H3K79me2 mark the intergenic enhancer, in future studies, it is worth investigating those enhancers’ regulatory impact on genes via long-range chromatin interaction by combining sequential ChIP with Hi-C assay to gain insight into why those enhancer elements are harbored at predominantly heterochromatic LB1 region.

Interestingly, we found that the G-rich promoters of the genes that are known to associate with and are regulated by zinc finger TFs are enriched in the genes at the LB1 compartment. G-rich DNA motifs are likely to form G-quadruplexes in different physiological conditions, and the promoters containing G-quadruplexes were shown to interact with zinc finger family TFs [78]. It is possible that the promoters associated with LB1 regions may contain G-quadruplexes and thus recruit zinc finger TFs to regulate gene expression [79] and this would also be an interesting mechanism to explore in future studies.

Nuclear periphery-localized genes are known to misregulate their expression in laminopathy diseases [80]. In addition, using mouse models, it has been shown that misregulation of LB1 in the nuclear periphery of striatal medium-sized spiny and CA1 hippocampal neurons is linked to Huntington-mediated neurodegeneration and therefore, the LB1-associated actively transcribing genes may bear a pathological relevance in neurons [81]. In our study, by compiling the LB1-associated transcriptionally active genes in OPCs with Clinvar data, we found that several candidates (e.g., Fkrp, Dcx) from our gene sets are associated with known neuronal disease-related dystrophies in humans (Fig. 4G), suggesting that LB1-associated genes may have clinical relevance in some laminopathy diseases.

In summary, we raised awareness to the presence of heterogeneous NPCs in retinoic acid-stimulated *in vitro* mESCs-derived NPC cultures, which may guide future investigations using *in vitro*-derived NPCs as a model system. We also present evidence that the LB1 compartment of mESCs and OPCs contains a significant number of actively transcribing genes that exhibit unique molecular features. Thus, we raised the importance of investigation of the heterochromatin and euchromatin organization in other cell types using as a model system to study the gene expression and chromatin architecture. Further, we showed that the transcriptionally active gene bodies at the LB1 region are enriched for G and C, and the promoters of those genes are G-rich and are known to interact mainly with zinc finger family TFs’. Our data support a model whereby the complexity and activity of genes associated with the LB1 compartment is cell-type specific and can change upon differentiation.

## Acknowledgements

We wish to thank members of the Spector lab for critical discussions and advice throughout this study. We thank Osama El Demerdash for assisting with setting up a computational pipeline. We would also like to thank Sara Goodwin (CSHL) for assisting with DNA sequencing. We acknowledge the CSHL Cancer Center Shared Resources (Microscopy and Next-Gen Sequencing) for services and technical expertise (NCI 2P3OCA45508). Sequencing analysis was performed using equipment purchased through NIH grant S10OD028632-01.

## Author Contributions

G.I.B. and D.L.S. conceived and designed the overall study. G.I.B. designed and performed the experiments and analyses. P.N. performed the IF experiments and T.W. assisted with imaging experiments. G.I.B. and D.L.S. wrote and edited the manuscript.

## Data availability

The RNA Pol II pSer2 ChIP-seq data (raw and processed) supporting this study’s findings have been deposited into the Gene Expression Omnibus (GEO) under accession code GSE246040. We have provided all other data supporting the findings of this study with the manuscript.

## Consent for publication

All authors agree with the publication of this article.

